# Avoidance learning reduces intricate covariation between boldness and foraging behavior in a generalist predator

**DOI:** 10.1101/2022.12.13.520202

**Authors:** Chi-Yun Kuo, Hao-En Chin, Yu-Che Wu

## Abstract

Many predators avoid unprofitable prey by learning to use visual features of the prey as reliable indicators of quality. Despite a rich literature on avoidance learning, we are still in the process of understanding how individuals might change their behavior towards unprofitable prey as information accumulates, the individual-level variation in avoidance foraging behavior, and whether learning would preserve or reduce this variation. In this study, we investigated how avoidance foraging behavior varied in a generalist lizard predator in relation to sex, population of origin, and boldness and tested the effects of learning on the variation in avoidance foraging behavior. We collected data on boldness and avoidance foraging behavior in individuals from two allopatric populations and compared variation in behavior at the beginning and end of learning experiments, in which individuals were presented with normal- and bitter-tasting prey that also differed consistently in color. We found that even though bitter prey elicited strong negative responses, lizards overall did not avoid consuming fewer such prey with learning. Instead, lizards learned to prioritize on palatable prey as they acquired more information. Our data revealed intricate covariation between boldness and avoidance foraging behavior, and the form of this covariation varied between populations, sex, and prey color. Learning overall reduced individual variation in foraging behavior, as well as the degree of covariation with boldness. Our findings highlighted the nuanced manners in which avoidance foraging behavior might manifest and suggested that learning could quickly mitigate fitness difference due to variation in behavior.

**Lay Summary:** Predators can learn to avoid unprofitable prey using certain visual cues or signals, but avoidance foraging behavior can manifest in more subtle ways than simply “to eat or not to eat”. We reported changes in foraging sequence and priority in a generalist lizard predator after learning and found that the nature of these changes depended on sex, population of origin, boldness, and prey color. We also found that learning made individuals more similar in behavior.

## Introduction

Recognizing and avoiding unprofitable prey reliably is an important task for all predators. Avoidance learning in this context describes adaptive changes in foraging behavior as predators gather information about prey profitability through repeated interactions (e.g., Speed 1993; Speed 2001; Sherratt 2011; Aubier and Sherratt 2020). Studies of avoidance foraging and learning have advanced our knowledge at the intersection of behavioral ecology and evolution, offering crucial insights on topics such as optimal foraging, the functioning of warning signals, and the evolution of Müllerian mimicry (Speed et al. 2000; Skelhorn and Rowe 2007; Barnett et al. 2012; Miller and Pawlik 2013; Kang et al. 2016; Páez et al. 2021). These studies indicate that predators are often able to base their foraging decisions on visual features of the prey when these features serve as reliable indicators of profitability.

Despite a rich body of literature, there remains three knowledge gaps in the study of avoidance foraging and learning. First, the vast majority of existing research on avoidance learning were performed on a small number of passerine birds (e.g., Skelhorn and Rowe 2006a; Halpin et al. 2008; Ihalainen et al. 2008; Rowland, Hoogesteger, et al. 2010; Smith et al. 2016; Rojas et al. 2019). Even though these studied have been important in establishing and testing theories and mechanisms of avoidance learning, we still only have a limited understanding of how predators in other taxa might modify their responses to unprofitable prey through learning. As a consequence, whether avoidance learning might manifest differently in other taxa remains a largely unexplored question. Second, existing studies on avoidance learning have almost exclusively focused on the overall level of avoidance towards unprofitable prey, calculated as the percentage of such prey not being attacked or consumed in choice experiments (Schall 2000; Long and Hay 2006; Webb et al. 2008; Rowland, Mappes, et al. 2010; Aronsson and Gamberale-Stille 2012; Taylor et al. 2016; Mulà et al. 2022). While the overall level of avoidance is a key metric of learning outcomes, predators might modify other aspects of avoidance foraging behavior, such as foraging latency and priority, towards unprofitable prey as they acquire more information. Yet, studies have rarely investigated this possibility, even when the focal predators exhibited strong negative responses towards unprofitable prey yet did not avoid consuming them (e.g., Skelhorn and Rowe 2009).

More importantly, individual-level variation in avoidance learning is often not examined in detail, despite the recognition that such variation plays an important role in animal behavior (Sih et al. 2004; Sih et al. 2015; Laskowski et al. 2022). A major source of individual variation in learning is differences in personality. The correlated manifestations of risk-taking tendency in exploratory and foraging contexts have been widely recognized as a behavioral syndrome or animal temperament; individuals that explore unfamiliar environments more readily tend to exhibit a lesser extent of foraging neophobia and higher level of foraging tolerance (Sih et al. 2004; Réale et al. 2007; Greggor et al. 2015). The correlation between personality and cognitive ability or learning style has also been increasingly investigated and debated over the past decade, which has led to the recognition that the nature and strength of such correlation often depends on the type of personality and learning traits being compared (Sih and Del Giudice 2012). It was initially hypothesized that there would be a positive correlation between exploration or boldness and learning ability as a result of environment-specific selection; more stable environments would favor fast explorers that are also fast learners, whereas more unpredictable environments would favor more cautious explorers that are slow but more flexible learners (Guillette et al. 2009; Carere and Locurto 2011; Sih and Del Giudice 2012; Griffin et al. 2015). However, a recent meta-analysis on 25 studies showed that even though exploration and boldness were correlated with learning ability, the direction of such correlation was equally likely to be positive or negative. That is, bolder and faster explorers tended to be either particularly fast or particularly slow learners (Dougherty and Guillette 2018). Moreover, there was extensive variation both within and across major groups, indicating that findings from better studied systems may have limited generality. Notably, only half of these 25 studies were from non-avian taxa; insects and non-avian reptiles were particularly understudied, represented by one study each. It was also noteworthy that the meta-analysis by Dougherty and Guillette (2018) included only one study that tested avoidance learning (Exnerova et al. 2010). Other factors, such as sex and the degree of allopatry, can also contribute to how boldness and learning styles might covary. The correlation between personality and learning ability was found to be stronger in males than in females (Dougherty and Guillette 2018). Even though population-specific effects have not been formally included in the meta-analysis of Dougherty and Guillette (2018), populations that are more spatially separated tends to differ in the strength of behavioral covariation in general (Garamszegi et al. 2012). Given the trait-specific nature of these findings, however, it still remains to be tested to what extent these results will hold for relationship between boldness and avoidance learning.

Filling these knowledge gaps would not only offer useful information on how individuals possessing different characteristics might avoid unprofitable prey in different ways, it will also enable us to answer an important question: does avoidance learning change foraging behavior while preserving individual-level variation, or does learning homogenize avoidance foraging behavior between individuals (Figure 1)? Since the effect of learning on behavior can be considered as a form of phenotypic plasticity in response to repeated exposure to the environment (Wright et al. 2022), the answer to this question has important implications to behavioral evolution. If learning alters behavior while preserving individual variation, it would imply that either the magnitude of avoidance response or the optimal avoidance foraging behavior depends on the values of behavioral covariates (e.g., sex, boldness, etc.). Conversely, elimination of individual variation through learning could mean that there is likely one singular optimal response, achievable by all individuals (e.g., complete avoidance to highly unprofitable prey, (Chouteau et al. 2019). At present, this question is mostly addressed in light of indirect social learning, which is changes in foraging behavior from observing the interactions between other individuals and prey. Social learning generally reduces the difference in avoidance foraging behavior between individuals with first-hand information about prey profitability (demonstrator) and those without (observer)(Salva et al. 2009; Hämäläinen et al. 2020; Mulà et al. 2022). The degree of variation in avoidance foraging behavior before and after direct learning, however, is rarely compared explicitly.

**Figure 1.**
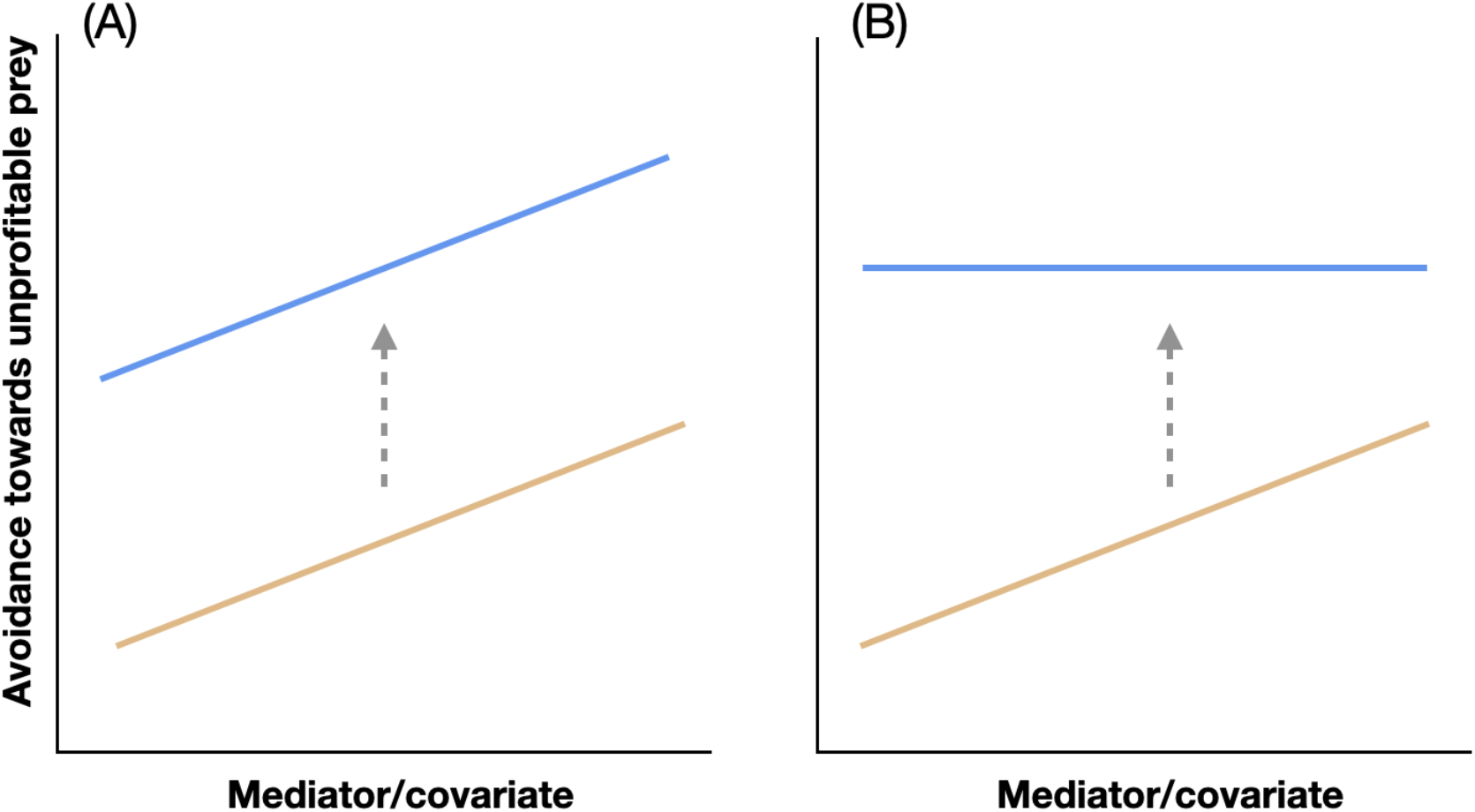
Conceptual diagrams showing the two competing hypotheses regarding how learning might affect the variation in avoidance foraging behavior in relation to another mediator variable or covariate. Brown lines are before learning, and blue lines are after learning. The y-axis represents values of some general avoidance measurement, not the overall level of avoidance specifically.

In this study, we quantified boldness in a generalist predator from two allopatric populations and repeatedly exposed them to prey that consistently differed in color (red vs. yellow) and taste (normal vs. bitter). We compared their foraging behavior between the beginning and the end of the experiment to answer the following questions: (1) is there a covariation between boldness and avoidance foraging behavior? (2) does the covariation between boldness and avoidance foraging behavior differ between sexes and populations? and (3) does the process of avoidance learning preserve or eliminate individual-level variation in foraging behavior? Based on existing knowledge, we predict that bolder individuals will exhibit lower avoidance towards unpalatable prey at the beginning of the experiment. We also predict that this relationship will be more prominent in males than in females. Even though the two focal populations were spatially separated, given the short time of divergence and a relatively small geographic scope of the study area, we did not have strong reasons to predict whether the two populations would differ in their foraging behavior and their responses to avoidance learning. Since the unprofitable prey in our study are merely unpalatable, we predict that avoidance learning will not eliminate individual variation in foraging behavior after learning.

## Materials and Methods

### Study species and husbandry

The common sun skink, *Eutropis multifasciata*, is native to south and southeastern Asian tropics and has a broad geographical distribution. This species was first recorded in Taiwan in 1992 and has quickly become invasive throughout central-to south-western part of the island (Ota et al. 1994; Chen et al. 2021). *Eutropis multifasciata* is an active forager with one of the most diverse diets among tropical skinks; observed food items included arthropods, snails, small vertebrates such as lizards and frogs, and plant materials (Ngo et al. 2014; Ngo et al. 2015). Their dietary composition can also vary markedly between seasons and regions (Ngo et al. 2014; Ngo et al. 2015). Avoidance learning and associated decision-making might be particular important for *E. multifasciata* than species with more specialized diets. We collected adults from two allopatric populations in southern Taiwan, Kaohsiung (KS: 22.64ºN, 120.26ºE) and Pingtung (PT: 22.48ºN, 120.57ºE), from March to August of 2022. We determined the developmental stage and sex of individuals based on external morphology and published mature snout-to-vent length (SVL)(male: 99 mm, female: 96 mm, (Sun et al. 2012). Lizards were brought back to the lab on the same day of capture. Each individual was housed separately (cage dimension: 30 × 20 × 25 cm) under a 12L:12D cycle and temperatures from 28 – 30ºC. Fresh water was provided daily, as well as crickets or meal worms as food except during avoidance learning experiment (see below).

### Boldness assays

We quantified boldness in the context of exploration and risk-taking in novel environments in *E. multifasciata*. Even though boldness and exploration were often treated as separate axes of animal personality (e.g., Réale et al. 2007), our study measured behaviors that related to both aspects (willingness to explore novel environments and responses to potential risk in the environment). We therefore used the term boldness to encompass our behavioral measurements. A pioneer dataset collected from a non-focal population showed that boldness remained the same for at least one month in captivity (Figure S1). We therefore assayed each individual twice, on the third and the seventh day after capture, to assess the repeatability of boldness. Boldness assays were performed in a terrarium (length × width × height: 90 × 30 × 45 cm) with beechwood chips bedding. An opaque division separated the terrarium into two areas (30 or 60 × 30 × 45 cm). Prior to the assay, we transferred a lizard from its home cage into the smaller chamber to acclimate for 10 mins. The smaller area contained a shelter and a rectangular brick from the lizard’s home cage, representing a more familiar and safer space, whereas the larger area was without any shelter or objects familiar to the lizard. We removed the division after the acclimation period and recorded the lizard’s behavior for 30 mins with a GoPro camera mounted one meter above the terrarium (Hero 8, GoPro, San Matteo, CA, USA). From each video, we extracted the following four variables: (1) moving latency, which was the time in second between the removal of the division to the time the lizard made the first directional movement, (2) latency to explore, which was the time in seconds before at least half of the lizard’s body moved into the larger area. If this never happened for a lizard, we assigned a value of 2000 to this variable, (3) number of movements per second, which was calculated as the number of directional movements of the lizard divided by the total observation time in seconds, (4) percent time spent moving, which was the amount of time in seconds that the lizard spent moving divided by the total observation time, (5) percent time being visible, which was the amount of time in seconds during which at least half of the lizard’s body was outside the shelter and not in the bedding divided by total observation time, and (6) percent time in the new environment, calculated as time in seconds during which at least half of the lizard’s body was in the large area divided by total observation time. We sprayed the bedding and the terrarium walls with 70% ethanol between trials to minimize the potential influence of residual chemicals from the previous lizard.

### Avoidance learning experiment

We quantified avoidance learning behavior of lizards after the second boldness assays were complete. The experiment consisted of five consecutive daily trials, during which lizards were exposed to prey that differed in color-taste combinations. This is a standard design for quantifying avoidance learning across taxa (e.g., Long and Hay 2006; Webb et al. 2008; Rowland, Wiley, et al. 2010; Taylor et al. 2014; Zvereva et al. 2018). Even though some studies performed a larger number of consecutive learning trials, changes in foraging behavior typically stabilized by the first three to five trials (e.g., Skelhorn and Rowe 2006b; Halpin et al. 2008; Aronsson and Gamberale-Stille 2012; Pegram et al. 2021). Lizards were fasted for two days prior to the first trial and were not given food other than experimental prey during the experiment. In each trial, a lizard was presented with four yellow crickets (painted with MONA Acrylic Colour S-209) and four red crickets (painted with MONA Acrylic Colour S-203) in its home cage. *Eutropis multifasciata* readily consumed normal-tasting crickets of both colors in preliminary observations. There were two treatment groups that differed in the color-taste combination of the prey. For the red group, we made red crickets bitter by injecting them with ∼ 0.3 ml aqueous solution of 0.07% denatonium benzoate (weight percentage). Denatonium benzoate was a nontoxic, colorless, odorless, yet extremely bitter chemical commonly used in avoidance learning experiments (Skelhorn and Rowe 2006a; Siddall and Marples 2008; Skelhorn et al. 2008). The concentration we use in our experiment were based on these previous studies and could elicit strong negative responses from lizards in preliminary observations, including initial prey rejection, wiping of the snout, and excessive secretion of saliva. The yellow crickets in this group served as palatable control and were injected with the same amount of water. The yellow group experienced the opposite color-taste combination. We randomly assigned lizards into each group while making sure the two groups were generally balanced regarding sex and population of origin.

We recorded the sequence of interactions between the focal lizard and the crickets for 15 mins or until the lizard attacked all crickets. We recorded a lizard’s interaction with a cricket as “attack” if it made a physical attempt to consume the cricket and as “consume” if the cricket was subsequently fully swallowed. An interaction labeled as “consume” was thus necessarily preceded by an “attack” interaction. We derived three variables describing a lizard’s avoidance foraging behavior: (1) foraging latency, recorded as the duration in seconds from the presentation of the crickets to when the lizard made the first attack (2) the type of first cricket attacked (control vs. bitter), (3) the percentage of bitter crickets among the first 50% crickets attacked (hereafter referred to as *foraging priority*); the lower the value, the more a lizard tended to prioritize on palatable cricket. We calculated this variable only if a lizard attacked at least four crickets, and (4) the percentage of bitter crickets consumed (hereafter referred to as *overall level of avoidance*), calculated as the number of bitter crickets consumed divided by the total number of crickets consumed. We removed all remaining crickets at the end of each trial.

### Statistical analyses

We performed all statistical analyses in R (version 4.2.2, R Core Team).

We assayed boldness in 45 adults from the two populations (11 females and 7 males from the KS population, 12 females and 15 males from the PT population). To confirm the repeatability of boldness and to compare boldness between sexes and populations, we first performed a principal component analysis (PCA) on moving latency, latency to explore, number of movements per second, precent time spent moving, percent time being visible, and percent time being in the new environment to reduce the dimension of data into a small number of principal components (PCs). We used the PC1 score of each lizard to represent its boldness (see Results). We built generalized linear mixed models (GLMM), with PC1 scores as the response variable, assay (first vs. second), sex, and population as fixed effect factors, and individual identity as a random effect factor. None of the predictors warranted exclusion based on variance inflation factor values (all values < 1.1). We did not observe any potential interactions between predictors from visual examination of the data and therefore did not include any interaction terms in the GLMM. No variance structure was specified in the initial GLMM. We followed the model validation recommendations using residual diagnostics in (Zuur et al. 2009). We detected unequal residual distribution between populations and modified the GLMM to allow populations to have different variances (the *varIdent* specification in the *gls* function). This updated model was superior to the original model and did not result in obvious heterogeneity (Akaike Information Criterion [AIC] score: 358 vs. 334). We interpreted the results using this updated model.

We used avoidance learning data from 38 of the 45 lizards with boldness data in subsequent analyses (20 male and 18 females), after excluding one boldness outlier and six other individuals that attacked fewer than four crickets in either the first or the last avoidance learning trials. We removed an additional 9 outliers with unusually long foraging latencies using Grubb’s test, but these individuals were not excluded in analyses when using other behaviors as response variables. Since changes in foraging behavior appeared to have stabilized by the fifth trial (Figure S2), for simplicity we used behavioral data from the first and the fifth avoidance learning trials to examine the effect of avoidance learning on the variation in foraging behavior. We initially included individual identity as a random effect factor, but doing so resulted in singular models with trivial estimated random effect variance (< 0.0001). We therefore constructed separate generalized linear models (GLM) for each of the four foraging behavior variables, with boldness from the second assay (see Results), sex, trial, treatment group, and population as fixed effect predictors. Variance inflation factor scores did not reveal severe multicollinearity among predictors (all values < 2).

For foraging latency, we performed zero-inflated gamma regressions using the *glmmTMB* package to account for a substantial presence of zeroes in the response variable (Brooks et al. 2017). The zero-inflated gamma regressions first tested whether any predictors were associated with a higher likelihood of zero foraging latency (the zero-inflated component) and then tested whether changes in predictors were related to variation in non-zero foraging latencies (the conditional component). As considering all two-way interactions resulted in an overfitted and non-convergent model, we only considered interactions between boldness and each of the other predictors as the starting model and used iterative model selection based on AIC that allowed backward elimination. For the type of first prey attacked, we performed Bernoulli regressions to account for the binary nature of the response variable. Foraging priority and overall level of avoidance were both proportions derived from count data, and we performed logistic regressions when using them as response variables. Due to presences of zeroes in foraging priority, we performed the complementary log-log transformation on the variable in logistic regression. Visual examination of the data indicated potential two- and three-way interactions between predictors. We therefore iteratively searched for the best model based on AIC, starting with a model with all two-way interactions and allowing both backward elimination and forward addition with the *step* function in the *stats* package. We allowed the inclusion of at most three-way interactions in the model selection. Once the best model was found, we first confirmed that there was no overdispersion (for logistic GLMs) and then validated the model by performing residual diagnostics following Zuur et al. (2009) using the *DHARMa* package (Hartig, 2016).

When presenting results, we followed the recommendation by Muff et al. (2021) and interpreted the P-values in the language of evidence instead of significance dichotomy. For results from the logistic GLMs, we reported Cohen’s D along with test statistics and P-values for individual factors using the *effectsize* package (Ben-Shachar et al. 2020) and interpreted their magnitudes according to (Gignac and Szodorai 2016). We also calculated McFadden’s pseudo-R square for these regressions to assess the goodness-of-fit of the best model (McFadden 1974). Since no standardized effect sizes for individual factors were appropriate for GLMM (Pek and Flora 2018; Rights and Sterba 2019), we calculated semi-partial R squares using the *r2glmm* package for evaluating the contributions of fixed effect factors instead (Jaeger et al. 2017). We were not aware of any standardized effect size measures for zero-inflated gamma regression and only reported test statistic and P-values.

## Results

### Boldness: repeatability and variation

The first principal component (PC) summarized 68% of the total variation from the six measured variables and represented overall boldness (Table S1). Individuals with higher PC scores were shyer; they waited longer to move and to enter the large area, and spent less time being visible, moving, and being in the new environment (Table S1). The data showed no evidence that boldness differed between the two assays, confirming that boldness was repeatable over the study period (Table S2, Figure 2A). We also found no evidence that boldness differed between the sexes (Table S2, Figure 2B). However, there was very strong evidence that the KS individuals were bolder than their PT counterparts (Table 1, Figure 2C); population alone explained 41% of the generalized variance.

**Figure 2.**
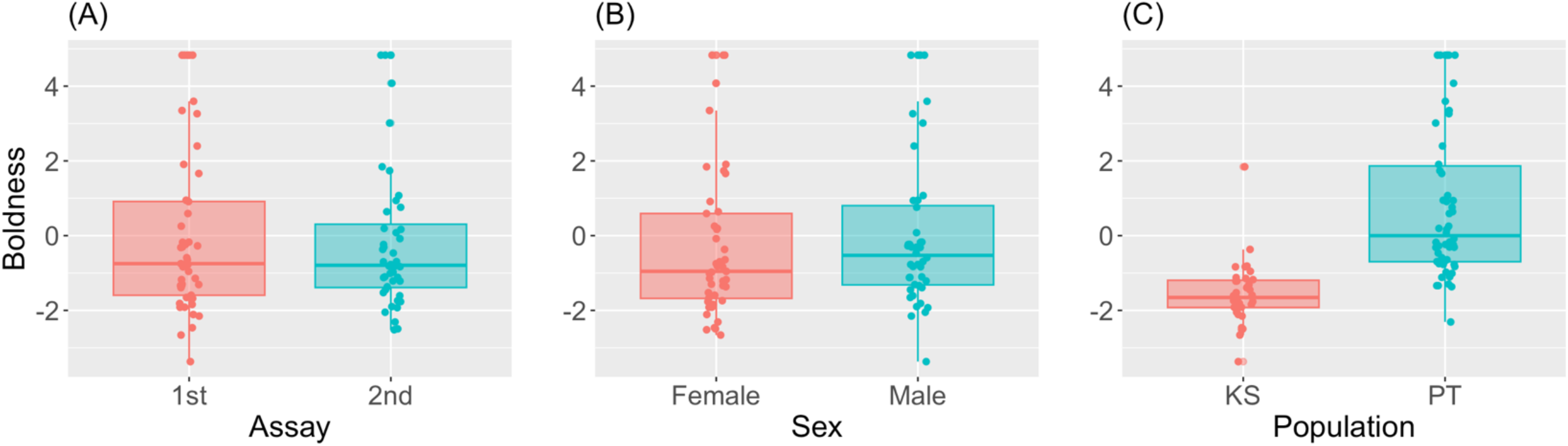
Repeatability of and variation in boldness in *E. multifasciata*. Lower values on the y-axis denote higher level of boldness. (A) Boldness values between the first and the second assay, confirming high repeatability. (B) Boldness between males and females, indicating no sexual difference. (C) Boldness between the two focal populations. The KS individuals were bolder than their PT counterparts.

**Table 1.** Summary of statistical analyses. Results were from GLMM for the analysis on the repeatability of and variation in boldness and GLM for analyses involving avoidance foraging behavior. Due to space limitation, only factors with P < 0.05 were presented.

| Factor | Coefficient | Statistic<sup>1</sup> | P | Evidence strength<sup>2</sup> | Cohen's D | Effect magnitude<sup>3</sup> |
| --- | --- | --- | --- | --- | --- | --- |
| Boldness (GLMM) |  |  |  |  |  |  |
| Population | 2.42 | 7.19 | < 0.0001 | Very strong | NA | NA |
| Foraging latency (zero-inflated gamma GLM): the conditional component |  |  |  |  |  |  |
| Sex | 0.63 | 2.04 | 0.041 | Moderate | NA | NA |
| Population | 1.56 | 3.75 | 0.0002 | Very strong | NA | NA |
| Boldness | 0.72 | 2.28 | 0.023 | Moderate | NA | NA |
| Sex:boldness | 0.68 | 2.25 | 0.024 | Moderate | NA | NA |
| Type of first prey attacked (GLM) |  |  |  |  |  |  |
| Sex:treatment | 3.52 | 2.17 | 0.030 | Moderate | 2.22 | Large |
| Sex:boldness | 7.44 | 2.06 | 0.040 | Moderate | 4.10 | Large |
| Treatment:trial | 1.17 | 2.11 | 0.035 | Moderate | 0.64 | Large |
| Treatment:boldness | 1.22 | 2.01 | 0.044 | Moderate | 0.67 | Large |
| Trial:population | 2.70 | 2.42 | 0.015 | Moderate | 1.49 | Large |
| Trial:boldness | 1.57 | 2.36 | 0.018 | Moderate | 0.86 | Large |
| Population:boldness | 4.46 | 2.21 | 0.027 | Moderate | 2.46 | Large |
| Trial:population:boldness | 1.86 | 2.59 | 0.010 | Strong | 1.03 | Large |
| Foraging priority (GLM) |  |  |  |  |  |  |
| Treatment | 0.70 | 3.15 | 0.002 | Strong | 0.38 | Small |
| Sex:boldness | 0.27 | 2.45 | 0.014 | Moderate | 0.15 | Very small |
| Treatment:boldness | 0.25 | 2.13 | 0.033 | Moderate | 0.18 | Very small |
| Trial:boldness | 0.33 | 2.08 | 0.038 | Moderate | 0.18 | Very small |
| Trial:population:boldness | 0.37 | 2.27 | 0.002 | Strong | 0.21 | Small |
| Overall level of avoidance (GLM) |  |  |  |  |  |  |
| Trial:sex:boldness | 0.12 | 2.31 | 0.021 | Moderate | 0.07 | Very small |

1. It was t-statistic for the GLMM and z-statistic for the GLM 2. Evidence strengths were based on Muff et al. (2021) 3. Effect size magnitudes were based on Gignac and Szodorai (2016)

### Learning-independent variation in avoidance foraging behavior

There was no evidence that any predictors were associated with more occurrences of zero foraging latencies (Table S3). However, there was very strong evidence that the PT individuals had longer foraging latencies (Table 1, Figure 3A). We also found moderate evidence for an interaction between sex and boldness; males had shorter foraging latencies compared to females, and non-zero foraging latency decreased with boldness more in males than in females (Table 1, Figure 3B). The best model with type of first prey attacked as the response variable explained 42% of the total variance. We found moderate evidence for sex-treatment and sex-boldness interactive effects (Table 1). Bolder lizards of both sexes in the yellow group tended to attack palatable crickets first more often than shyer individuals. This covariation was weaker in females and slightly reversed in direction in males in the red group (Figure 3C, D). Regarding foraging priority, the best model explained 27% of the total variance. There was strong evidence for a small treatment effect and moderate evidence for a very small treatment-boldness interaction (Table 1). Individuals in the red group attacked fewer bitter crickets among the first 50% crickets attacked than their counterparts in the yellow group (Table 1, Figure 3E). In addition, bolder individuals in the yellow group tended to prioritize on normal crickets, whereas such foraging priority was nonexistent among lizards in the red group (Table 1, Figure 3E). Finally, we found moderate evidence of a very small sex-boldness interactive effect (Table 1). Bolder females had a slight tendency to prioritize on control crickets, but this trend was absent in males (Table 1, Figure 3F). The best model for overall level of avoidance explained 14% of total variance. However, the data did not show evidence for any learning-independent variation in the percentage of bitter crickets consumed (Table S5).

**Figure 3.**
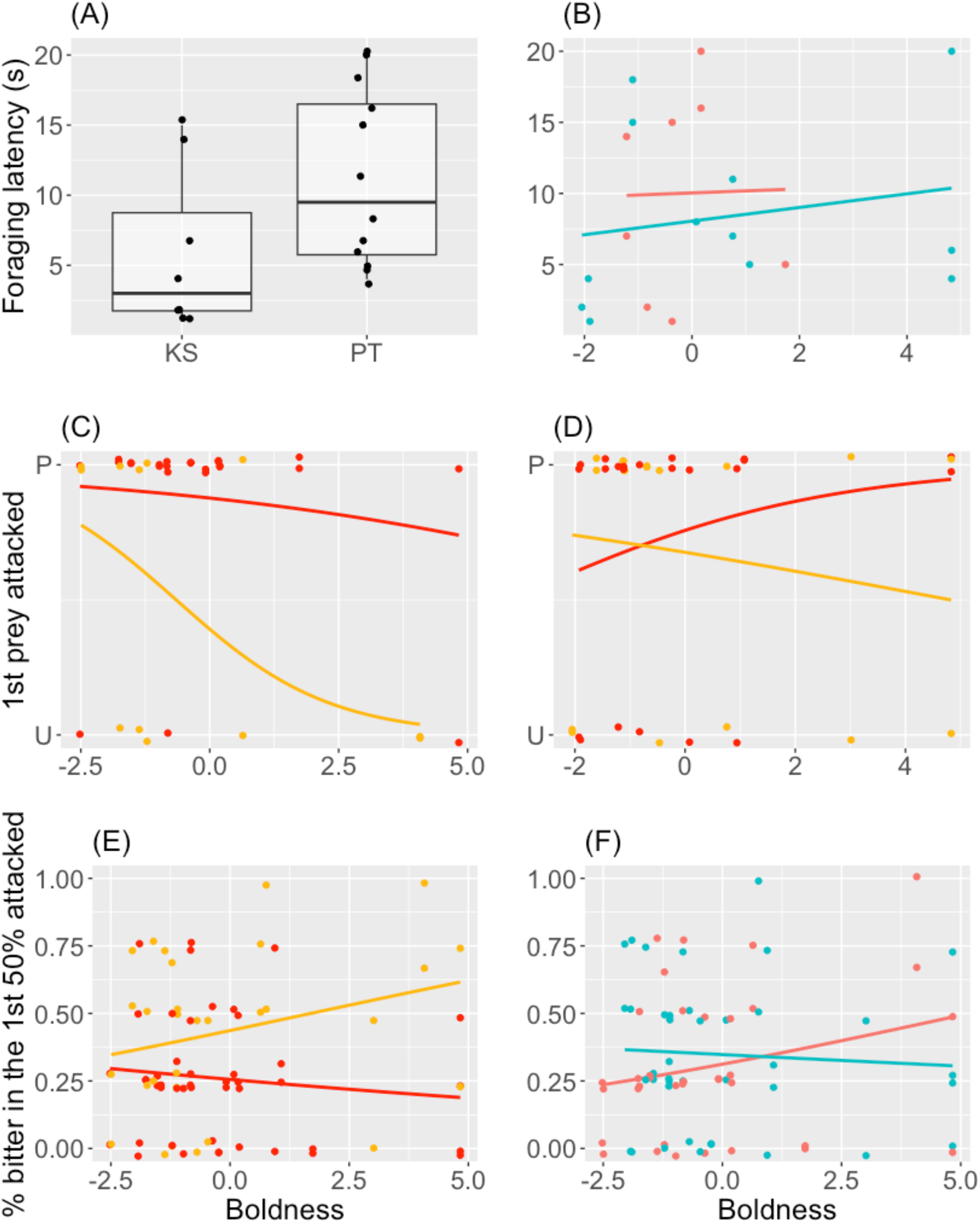
Variation in avoidance foraging behavior independent of learning (A) Foraging latencies differed between the two populations. (B) Sexual difference in the pattern of covariation between boldness and foraging latency. The type of first prey attacked (U: unpalatable, P: palatable) covaried with boldness in different ways between the two treatment groups (red and yellow) in females (C) and males (D). Foraging priority covaried with boldness differently in individuals belonging to the two treatment groups (E) and the two sexes (F, red dots and line: females; blue dots and line: males).

### Effect of avoidance learning on the variation in foraging behavior

Generally, lizards tended to attack control prey more often at the end of the experiment (Figure 4A-D). There was strong evidence for a large interactive effect between trial, population, and boldness on the type of first prey attacked (Table 1). In the KS population, bolder individuals were more willing to attack bitter prey first at the beginning of avoidance learning. However, it was shyer individuals from the PT population that showed the same tendency during the first trial. The process of avoidance learning eliminated this population-level difference; the slopes of the logistic regressions were much flatter and less varied between the two populations at the end of the experiment (Figure 4A, B). There was also moderate evidence indicating large treatment-trial and treatment-boldness interactive effects (Table 1); bolder individuals in the yellow group tended to attack palatable crickets first in the first trial, whereas this tendency was not present in individuals in the red group (Figure 4C, D). With respect to foraging priority, lizards prioritized more on control crickets at the end of the experiment (Figure 4E, F). There was strong evidence of a small interactive effect between trial, population, and boldness (Table 1); bolder lizards from the KS population and shyer lizards from the PT population both had lower tendencies to prioritize on palatable crickets in the first trial. This covariation was reversed in the KS individuals and largely absent in the PT individuals at the end of avoidance learning (Figure 4E, F). Even though the bitter taste of the experimental crickets induced strong negative responses, *E. multifasciata* did not avoided consuming more bitter crickets as the experiment progressed (Table S5). The proportion of bitter crickets consumed decreased only slightly from 39% to 35% between the first and the last trials. The only predictive factor we had nontrivial evidence for was a very small interactive effect between sex, trial, and boldness (Table 1). Bolder females and shyer males both consumed fewer bitter crickets at the beginning of the experiment, but the trend was reversed in both sexes in the last trial (Figure 4G, H).

**Figure 4.**
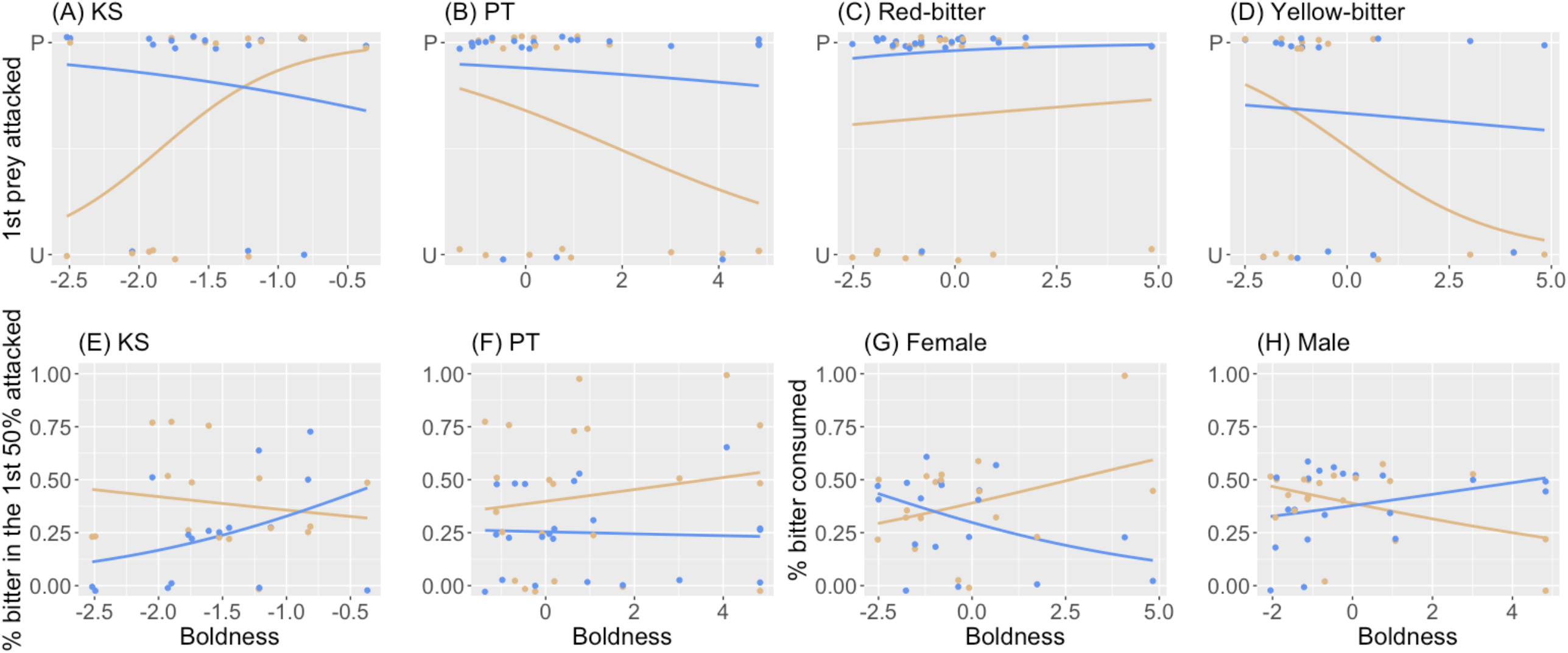
Learning-induced changes in the covariation between boldness and avoidance foraging behavior. (A-D) Type of first prey attacked (U: unpalatable, P: palatable). (E, F) Foraging priority. (G, H) Overall level of avoidance. Brown dots and lines are results from the first learning trial, and blue dots and lines are those from the fifth trial. (A), (B), (E), and (F) show difference between populations (KS vs. PT). (C) and (D) are difference between treatment groups. (G) and (H) are difference between the sexes.

## Discussions

### Intricate variation in avoidance foraging behavior independent of learning

Our results revealed complex variation in avoidance foraging behavior independent of learning. Most of the variation occurred in the form of different patterns of covariation with boldness between sexes, populations, and the color-taste combination of the prey. In general, we found support for the prediction that bolder individuals would be less aversive to bitter prey. Consistent with the predictions, the KS individuals, which were much bolder than their PT counterpart, were overall quicker to attack (Figure 3A). Bolder individuals from the KS population were also more prone to attacking unpalatable crickets as the first target during the first trial (Figure 4A). Bolder males, but not females, also had shorter foraging latency (Figure 3B). The prediction that the covariation between boldness and avoidance foraging behavior would be stronger in males was better supported when bitter prey were associated with red color. Bolder males in the red group tended to attack bitter crickets as the first target more often, and bolder individuals of both sexes in the red group had a slightly lower tendency to prioritize on control crickets (Figure 3D, E). Data from females were partially supportive of the prediction. The covariation between boldness and foraging latency was almost absent in females regarding foraging latency and overall level of avoidance. Interestingly, we found that bolder females tended to attack palatable crickets first throughout the experiment, especially when prey unpalatability was associated with yellow color (Figure 3C).

The prediction concerning the nature of boldness-avoidance foraging covariation was overall not supported when bitter taste was associated with yellow color, and this was largely due to contrasting behavior of shyer individuals in the two treatment groups (Figures 3C-E). Regardless of the sex, it was shyer individuals that tended to target bitter crickets first, and this trend was even more prominent in females (Figure 3C, D). Shyer individuals in the yellow group also prioritized on palatable crickets to a lesser extent (Figures 3E). To our knowledge, this is the first report of different color-taste combination resulting in contrasting patterns of covariation between boldness and avoidance foraging behavior. It was unlikely that shyer individuals had higher bitter tolerance or were less choosy foragers overall, because this trend was not present in the red group. Instead, we hypothesize that this observation was the combination of the color yellow being less effective in facilitating the link with prey unpalatability and shyer individuals being slower learners. In the red group, the higher efficacy of the color red might have made the difference in learning ability less prominent. Indeed, previous studies have reported that colors differ in their efficacy as avoidance learning cues when overall level of avoidance is concerned (Skelhorn and Rowe 2006a; Taylor et al. 2014; Taylor et al. 2016), and our results might have been a manifestation of this fact in different aspects of avoidance foraging behavior.

Another notable contradiction to the prediction was that the PT individuals exhibited the reverse from what we had predicted (Figure 4B). However, it was possible that data from the two populations revealed a nonlinear pattern in which the tendency to attack palatable crickets as the first target was highest in individuals with intermediate boldness (Figures 4A, B). With the data at hand, a GLM with both boldness and population as predictors was superior to one with a second-degree polynomial of boldness but not population as predictor (AIC: 84 vs. 86). This represented an intriguing hypothesis to be further tested with data from more individuals and more populations.

### Avoidance learning reduces variation in foraging behavior

There were prominent, contrasting covariation between boldness and the type of first prey attacked during the first trial with respect to population and treatment group (Figures 4A-D). These strong tendencies observed from the first trial were curious in themselves, since the lizards should have had no information about the color-taste association to base their decisions on. One possible explanation is that individuals were able to assess prey palatability through tongue-flicking, a behavior that scincid lizards use to collect olfactory information from the surrounding air. Given the fact that the bitter chemical was odorless and injected inside the crickets rather than applied externally, we suspected that the information about prey palatability gathered through tongue-flicking was likely much more limited than other sampling behavior, such as taste rejection in birds (e.g., Skelhorn and Rowe 2006c; Skelhorn and Rowe 2006d; He et al. 2022).

We found that learning did not make *E. multifasciata* consume fewer bitter crickets but changed their foraging sequence and priority; with more information on the color-taste coupling of the prey, *E. multifasciata* targeted and consumed palatable prey with higher priorities, regardless of their boldness (Figures 4A-F, brown vs. blue curves). This underscored the flexible and opportunistic foraging habits of *E. multifasciata*, which was commonly observed in other generalist predators (e.g., Terraube and Arroyo 2011; Diller et al. 2019). In the field, the presence of high-quality prey is likely staggered and unpredictable. By prioritizing on palatable crickets first, *E. multifasciata* might be able to avoid consuming bitter crickets altogether if more palatable prey appeared later in time. Since we did not offer additional prey during the experiment, the need for nutrition likely outweighed the negative foraging experience from prey bitterness, hence the low overall level of avoidance. Our data could not concretely validate this hypothesis, but it was clear that individuals by and large converged toward the same foraging behavior by the end of the experiment. Our findings therefore were in line with the prediction that learning not only changed but also homogenized behavior (Figure 1B), instead of preserving individual-level variation despite behavioral changes (Figure 1A).

The evolutionary implication of our findings is that any difference in fitness as a consequence of boldness-avoidance foraging covariation can potentially be quickly and largely mitigated by learning. This learning-based plasticity could allow variation in boldness and other boldness-associated behavior to persist among individuals and manifest in other contexts. It would be intriguing to examine how quickly the variation in avoidance foraging behavior re-emerges as the connection between color and prey quality deteriorates due to forgetting and how much more quickly the foraging behavior changes when individuals are re-exposed to the same color-taste coupling of the prey.

### Conclusions and future directions

In this study, we presented useful information on avoidance foraging behavior outside a small number of avian species and highlighted aspects of foraging behavior that were rarely examined in other avoidance learning studies. We found complex variation in avoidance foraging behavior in a generalist lizard, and nature of variation depended on sex, population of origin, boldness, and the exact color-taste combination of the experimental prey. Our results showed that avoidance learning changed the foraging sequence and priority without noticeable effects on the level of prey avoidance. Overall, avoidance learning eliminated both individual- and population-level variation in foraging behavior.

We would like to end by offering a few directions for future research on behavioral covariation involving boldness, avoidance foraging, and learning. First, what is the role of genetics versus plasticity in determining the pattern and degree of covariation between boldness and avoidance foraging behavior? Even though a large number of studies have estimated the heritability of numerous behaviors individually (reviewed in Dochtermann et al. 2019), we are still in the process of uncovering the degree of heritability for behavioral covariation (Rudin et al. 2019). The answer to this question is likely trait- and taxa-dependent. A recent study on a lizard revealed that exploratory behavior and spatial cognition were both highly repeatable yet showed no heritability (Meester et al. 2022). The generality of their findings merits further testing. Second, it would be informative to test whether the population-level difference in the boldness-foraging behavior covariation revealed by our data actually reflected different portions of the same non-linear relationship between the two behaviors. Data from more populations would help address this question, depending on whether such data revealed more population-specific patterns of covariation or strengthen the same parabolic relationship shown in our current data. Lastly, the time frame of potential behavioral reversal and the re-emergence of behavioral covariation due to forgetting is rarely quantified even in better studied avian systems but would offer valuable insights into how learning-associated plasticity might affect fitness (Wright et al. 2022).

## Funding

This work was supported by funding to CYK from the Taiwan National Science and Technology Council (grant number 110-2621-B-037-003) and Kaohsiung Medical University Research Foundation (grant number KMU-Q111001). All experimental procedures were approved by the IACUC at Kaohsiung Medical University (protocol number 110051).

We thank Hsiang-An Chao, I-Ting Hung, and Miao-Chu Tseng for their assistance in video analysis.

## Author contribution

CYK designed the study. CYK, HEC, and YCW collected lizards from the field, participated in animal husbandry, performed boldness assays and avoidance learning trials, and analyzed video recordings. CYK performed the statistical analyses and wrote the manuscript. All authors approved the submission of the manuscript. The raw data and the analytical R code can be accessed at https://github.com/chiyun0529/boldness-avoidance-foraging.

