## Supplemental Figures and Tables for "Avoidance learning reduces intricate covariation between boldness and foraging behavior in a generalist predator"

Table S1. Factor loading for the first two principal components of boldness and cumulative proportion of variance explained

| Variables | PC 1 | PC 2 |
| --- | --- | --- |
| Moving latency | 0.42 | -0.13 |
| Latency to explore | 0.45 | 0.003 |
| % time being visible | -0.38 | -0.53 |
| Movements per second | -0.40 | 0.41 |
| % time spent moving | -0.40 | 0.49 |
| % time in new environment | -0.40 | -0.54 |
| Cumulative % variance | 0.68 | 0.80 |

Table S2. GLMM results examining the repeatability of and variation in boldness.

| Factor | Coefficient | SE | df | t | P |
| --- | --- | --- | --- | --- | --- |
| Intercept | -1.63 | 0.22 | 43 | -7.30 | < 0.0001 |
| Assay (2^nd^) | 0.29 | 0.22 | 43 | 1.28 | 0.21 |
| Population (PT) | 2.42 | 0.34 | 42 | 7.19 | < 0.0001 |
| Sex (m) | -0.17 | 0.29 | 42 | -0.58 | 0.56 |

Table S3. Zero-inflated gamma regression results examining the variation in foraging latency

| Parameter | Coefficient | SE | z | P |
| --- | --- | --- | --- | --- |
| Conditional component | | | | |
| Intercept | 1.46 | 0.30 | 4.90 | < 0.0001 |
| Sex (m) | -0.63 | 0.31 | -2.04 | 0.041 |
| Population (PT) | 1.56 | 0.42 | 3.75 | 0.0002 |
| Boldness | -0.72 | 0.32 | -2.28 | 0.023 |
| Sex:boldness | 0.68 | 0.30 | 2.25 | 0.024 |
| Zero-inflated component | | | | |
| Intercept | 0.83 | 0.56 | 1.48 | 0.138 |
| Sex (m) | -0.03 | 0.61 | -0.05 | 0.956 |
| Population (PT) | -0.38 | 0.82 | -0.46 | 0.643 |
| Boldness | 0.01 | 0.27 | 0.03 | 0.973 |
| Sex:boldness | -0.17 | 0.28 | -0.59 | 0.553 |

Table S4. Bernoulli regression results examining the variation in the type of first prey attacked

| Parameter | Coefficient | SE | z | P |
| --- | --- | --- | --- | --- |
| Intercept | 5.94 | 3.49 | 1.70 | 0.089 |
| Sex (m) | 9.01 | 5.54 | 1.63 | 0.104 |
| Group (yellow) | -1.55 | 1.54 | -1.01 | 0.314 |
| Trial (5^th^) | -1.44 | 0.90 | -1.60 | 0.110 |
| Population (PT) | -4.81 | 3.62 | -1.33 | 0.184 |
| Boldness | 3.48 | 1.90 | 1.83 | 0.067 |
| Sex:group | 3.52 | 1.62 | 2.17 | 0.030 |
| Sex:population | -11.27 | 5.84 | -1.93 | 0.054 |
| Sex:boldness | 7.44 | 3.61 | 2.06 | 0.040 |
| Group:trial | -1.17 | 0.55 | -2.11 | 0.035 |
| Group:boldness | -1.22 | 0.61 | -2.01 | 0.044 |
| Trial:population | 2.70 | 1.12 | 2.42 | 0.015 |
| Trial:boldness | -1.57 | 0.67 | -2.36 | 0.018 |
| Population:boldness | -4.46 | 2.02 | -2.21 | 0.027 |
| Trial:population:boldness | 1.86 | 0.72 | 2.59 | 0.010 |
| Sex:population:boldness | -6.48 | 3.60 | -1.80 | 0.072 |

Table S5. Logistic regression results examining the variation in foraging priority

| Parameter | Coefficient | SE | z | P |
| --- | --- | --- | --- | --- |
| Intercept | -1.68 | 0.83 | -2.03 | 0.042 |
| Sex (m) | 0.01 | 0.22 | 0.04 | 0.970 |
| Group (yellow) | 0.70 | 0.22 | 3.15 | 0.002 |
| Trial (5^th^) | 0.36 | 0.25 | 1.48 | 0.139 |
| Population (PT) | 0.79 | 0.86 | 0.92 | 0.357 |
| Boldness | -0.53 | 0.50 | -1.08 | 0.281 |
| Sex:boldness | -0.27 | 0.11 | -2.45 | 0.014 |
| Group:boldness | 0.25 | 0.12 | 2.13 | 0.033 |
| Trial:population | -0.49 | 0.26 | -1.91 | 0.056 |
| Trial:boldness | 0.33 | 0.16 | 2.08 | 0.038 |
| Population:boldness | 0.69 | 0.51 | 1.36 | 0.175 |
| Trial:population:boldness | -0.37 | 0.17 | -2.27 | 0.023 |

Table S6. Logistic regression results examining the variation in overall level of avoidance

| Parameter | Coefficient | SE | z | P |
| --- | --- | --- | --- | --- |
| Intercept | -0.54 | 0.28 | -1.93 | 0.054 |
| Sex (m) | 0.20 | 0.38 | 0.53 | 0.595 |
| Group (yellow) | 0.55 | 0.30 | 1.82 | 0.069 |
| Trial (5^th^) | -0.07 | 0.08 | -0.91 | 0.365 |
| Boldness | 0.16 | 0.14 | 1.19 | 0.233 |
| Sex:group | -0.56 | 0.39 | -1.43 | 0.152 |
| Sex:trial | 0.06 | 0.10 | 0.64 | 0.521 |
| Sex:boldness | -0.34 | 0.19 | -1.81 | 0.070 |
| Trial:boldness | -0.07 | 0.04 | -1.74 | 0.083 |
| Sex:trial:boldness | 0.12 | 0.05 | 2.31 | 0.021 |

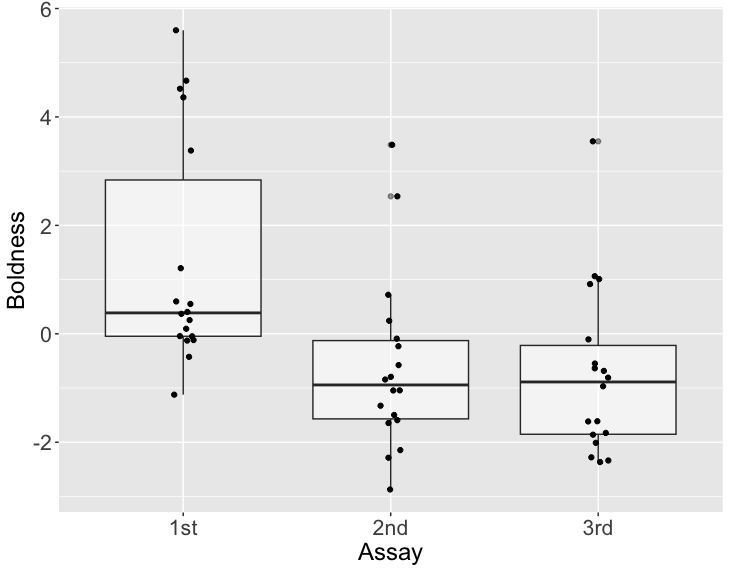

Figure S1. Pioneer dataset on the repeatability of boldness, collected from non-focal populations of *E. multifasciata.* The first, second, and third assays were conducted on the third, seventh, and the 30^th^ day after the individuals were brought back to the lab.

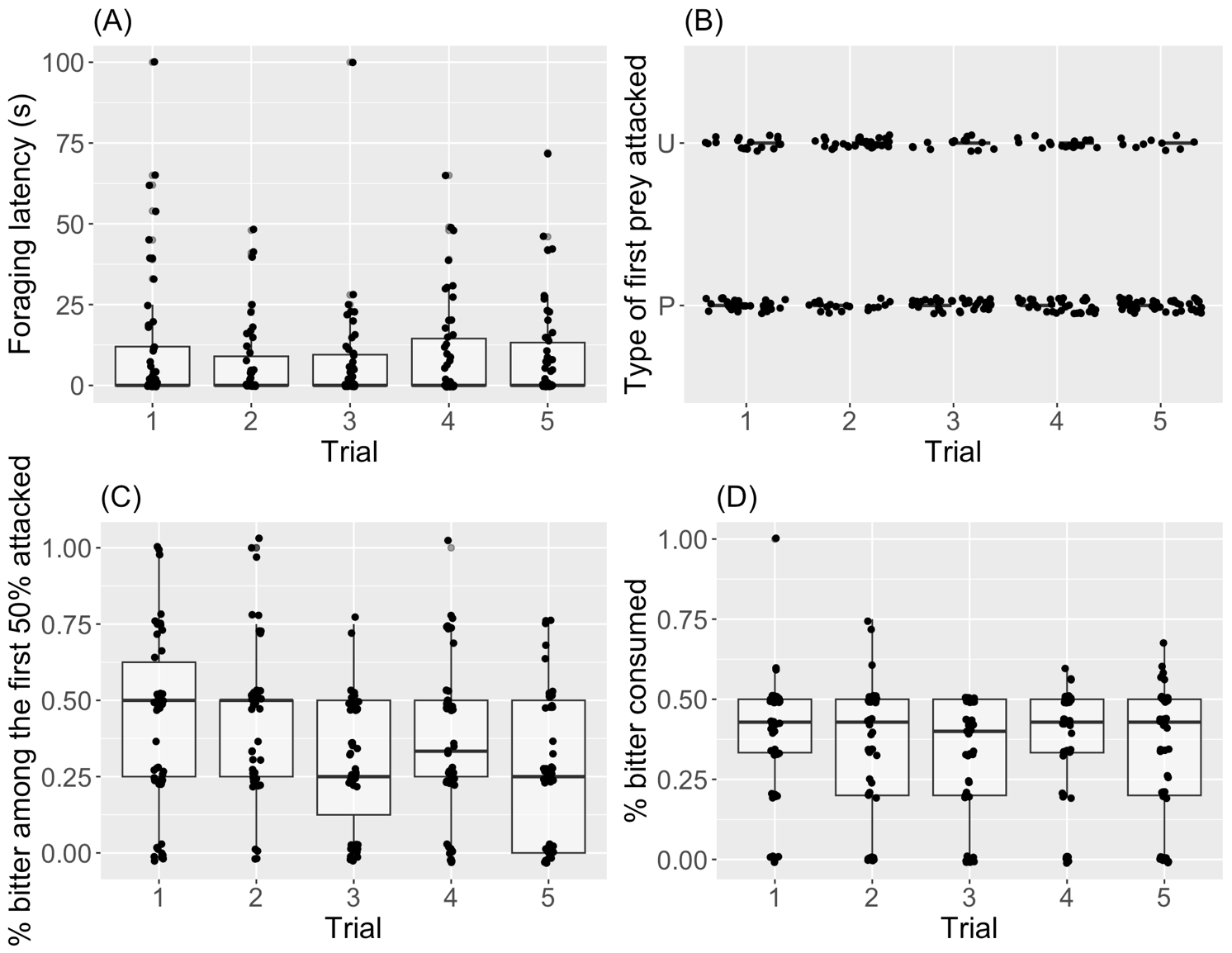

Figure S2. Changes in avoidance foraging behavior through the trials. (A) Foraging latency. (B) The type of first prey attacked (U: unpalatable, P: palatable). (C) Foraging priority. (D) Overall level of avoidance.
